# Evidence for allocentric boundary and goal direction information in the human entorhinal cortex and subiculum

**DOI:** 10.1101/466789

**Authors:** J. P. Shine, J. P. Valdés-Herrera, C. Tempelmann, T. Wolbers

**Author notes:** Equal author contributions (J. P. Shine, J. P. Valdés-Herrera). Corresponding Author: Dr. Jonathan Shine German Center for Neurodegenerative Diseases (DZNE) Leipziger Str. 44, 39120 Magdeburg.

## Abstract

In rodents, cells in the medial entorhinal cortex (EC) and subiculum code for the allocentric direction to environment boundaries, which is an important prerequisite for accurate positional coding. Although in humans boundary-related signals have been reported, there is no evidence that they contain allocentric direction information. Furthermore, it has not been possible to separate boundary versus goal direction signals in the EC/subiculum. To address these important questions, participants learned a virtual environment containing four unique boundaries, after which they underwent fMRI scanning where they made judgments about the allocentric direction of a cue object. Using multivariate decoding, we found information regarding allocentric boundary direction in posterior EC and subiculum, whereas in anterior EC and subiculum we could decode allocentric goal direction. These data provide the first evidence of allocentric boundary coding in humans, and are consistent with recent conceptualisations of a division of labour within the EC.

## Background

The entorhinal cortex (EC) provides the primary cortical input to the hippocampus^1^. Given its distinct profile of anatomical connectivity, different subregions of the EC have been hypothesized to convey different types of information that is combined in service of episodic memory and spatial navigation^2,3^. Specifically, in rodents, the medial EC (MEC) receives projections from regions processing spatial information, including the postrhinal cortex (primate parahippocampal cortex; PHC) and subiculum, whereas the rodent lateral EC (LEC) receives information from the object-sensitive perirhinal cortex. This pattern of anatomical connectivity has been shown to be preserved in humans, with posterior EC (homologous with rodent MEC) showing preferential connectivity with PHC^4^ and posterior subiculum^5^, whereas the anterior EC (homologous with rodent LEC) shares preferential connectivity with the perirhinal cortex^4^ and anterior subiculum^5^. These posterior and anterior divisions of EC and subiculum, therefore, have been hypothesized to code for “where" versus “what" information, respectively^6,7^. Recent evidence of spatial coding in the rodent LEC, however, has led to modifications of this model in which the MEC is proposed to support neural populations involved in spatial navigation (e.g., grid cells), whereas the LEC codes for external sensory inputs (e.g., landmarks or prominent objects in the environment)^8^.

In rodents, the MEC contains a number of different spatially-tuned neural populations, including border cells^9^. The coding of an environment’s boundaries is essential for neural computations that help to determine one’s spatial location. For example, path integration (i.e., the ability to update one’s spatial position on the basis of self-motion cues) is a process that invariably accumulates error^10,11^, and environmental cues, such as boundaries, help correct these noisy positional estimates^12^. Moreover, grid cell firing fields are anchored to the walls of an enclosure^13^, with irregular-shaped enclosures resulting in deformations of the grid cell’s characteristic hexagonal symmetry^14,15^. In the rodent hippocampus, the removal of environment boundaries leads to the degradation of place cells^16^, while expanding the size of a familiar environment by moving its boundaries leads to the commensurate expansion of a place cell’s firing field^17^. Cells coding for environment boundaries have been identified also in the rodent dorsal subiculum, with these so-called boundary vector cells containing information regarding not only the allocentric direction, but also the distance^18^, to a boundary. Accordingly, place cell activity has been modelled as the summed and thresholded input of these spatial properties describing an environment’s boundary position^19^. In sum, boundary coding is a fundamental component of spatial navigation.

In humans, boundaries have been shown to be behaviourally salient^20^,and recordings from the EC in intracranial patients have revealed increased theta frequency activity associated with the participant’s proximity to the environment walls during navigation in a virtual environment^21^. Similarly, boundaries have been shown to engage the hippocampus^22^, with univariate increases in hippocampal activity associated with the number of boundary elements to-be-imagined^23^. A recent study using intracranial recordings implicated the subiculum as the locus of this boundary coding effect^24^, again with reports of increased theta frequency activity in this region associated with the encoding of object locations proximal to a virtual environment’s walls. Despite evidence of boundary-related signals in the EC/hippocampus, these studies do not demonstrate that this neural activity contains the allocentric directional information necessary to support accurate positional coding^19^. Indirect evidence of allocentric boundary coding in the EC/subiculum has been found in a recent fMRI study using multivariate analysis methods, in which an allocentric goal direction signal, thought to reflect the simulated representations of head direction, was identified^25^. In this study, however, the heading direction to-be-imagined was determined by goal objects that were arranged in front of boundaries in the enclosed virtual environment. Consequently, it was not possible to fully disentangle whether this signal reflected allocentric goal or allocentric boundary direction. Moreover, this study lacked the anatomical resolution to differentiate subregions of the EC and subiculum. An outstanding question in the field, therefore, is whether there is evidence of an allocentric boundary direction signal in the human EC and subiculum.

In contrast to the MEC, neurons in the rodent LEC show far less spatial tuning. LEC neurons have been shown to code for space in the presence of objects, and there is evidence of trace memory for the previous location of a prominent object in the environment^26^. It has been hypothesised, therefore, that the function of the LEC may be to code for the positions of objects in space, which would constitute a landmark, or goal location^8^. In humans, a preference for object versus scene/spatial manipulations has been demonstrated in the anterior EC^6,7^, and is thought to reflect the region’s strong connections with object-specific perirhinal cortex. It is not clear, however, whether the human anterior EC is engaged also during spatial judgments regarding the allocentric direction of landmarks in the environment (i.e., object in place coding).

In the current study, we investigated the neural correlates of allocentric boundary and allocentric goal direction coding in the human medial temporal lobe, using a combination of immersive virtual reality and high-resolution fMRI. Importantly, we used anatomically informed masks of the EC and subiculum, and orthogonalised the relative contributions of allocentric boundary and allocentric goal processing. Given the different connectivity profiles, and proposed division of labour within the EC and subiculum, we separated our regions of interest (ROIs) into anterior and posterior portions to examine whether there was evidence of a longitudinal difference in decoding accuracy according to task. Specifically, we wanted to test whether posterior EC and subiculum were more involved in allocentric boundary coding, whereas anterior EC and subiculum coded information relating to the allocentric goal.

## Methods

### Participants

31 right-handed, young healthy adults (13 female; mean age 26.12 years, range = 20 – 33 years) participated in the experiment and were paid 31 euros for their time. All participants provided informed consent, and the experiment received approval from the Ethics Committee of the University of Magdeburg.

### General procedure

The experiment comprised two separate days of testing. On the first day, the participant learned the layout of a virtual environment (VE) outside of the scanner. On the following day of testing, the participant underwent high-resolution fMRI scanning.

### The virtual environment

The VE was created using WorldViz Vizard 5.1 Virtual Reality Software (WorldViz LLC, http://www.worldviz.com). It comprised a large grass plain (600 × 600 virtual metres^2^; invisible walls that prevented the participant from leaving the VE resulted in a 500 × 500 virtual metres^2^ explorable area) surrounded by four distinct global landmark cues, and contained four rectangular boundaries (2 × 4 × 40 virtual metres), each with a unique brick texture (Figure 1A). The global landmark cues (a mountain, a tower, a cathedral, and a city) were rendered at infinity, and indicated cardinal heading directions in the environment. During the experiment, however, the landmarks were never referred to using cardinal directions (i.e., north, south, east and west). In the VE, two of the four boundaries were arranged with their long axes spanning north-to-south, whereas the other two boundaries’ spanned east-to-west.

**Figure 1.**
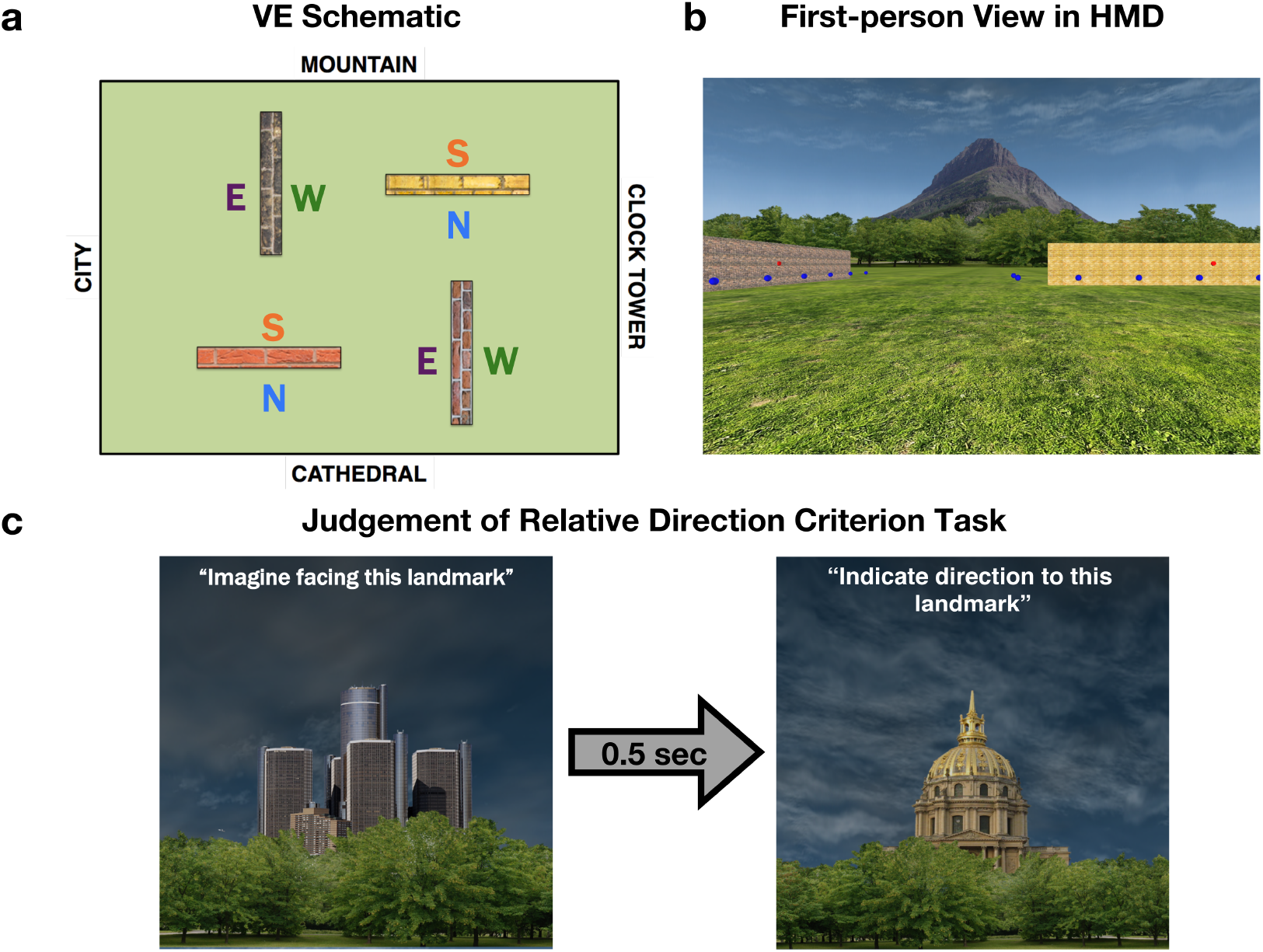
Schematic of the VE and criterion task during learning phase. (a) The VE comprised four boundaries, each with a unique texture. The long axis of two of the boundaries spanned North-South, and two spanned East-West. The VE was surrounded by four global landmarks rendered at infinity that provided information regarding cardinal direction in the environment. Each side of the boundary created an allocentric boundary either to the North, South, East or West. (b) In the learning phase the participant wore an HMD and controlled their orientation by physically rotating on the spot whereas translations were controlled via a button-press. During exploration, the participant was required to collect all blue tokens and activate all sensors located on the different sides of the boundaries. (c) After exploring the environment, the participant completed a JRD task in which they were presented with one of the global landmarks and asked to indicate the relative direction to another global landmark; participants were allowed to move on to the scanned test phase only after answering all JRD questions correctly.

### Training outside of the scanner

A head mounted display (HMD; Oculus Rift Development Kit 2) was used during training to provide an immersive learning experience. The participant stood during the experiment and was required to physically turn on the spot to change facing direction in the VE; translations were controlled via a button press on a three-button wireless mouse held in the participant’s right hand throughout the training phase. To promote exploration of the VE and its boundaries, the participant was required to collect blue tokens (0.25 virtual metre radius spheres positioned at a height of 0.8 virtual metres; the first-person view in the VE was rendered at 1.8 virtual metres) that formed a path around the four boundaries (Figure 1B). The participant was required to walk through each token after which it disappeared; the participant was free to collect the tokens in any order. Furthermore, to ensure that the participant was aware that the boundary was impassable, they were required to ‘activate’ red sensors located on each side of a boundary (eight sensors in total, positioned at a height of 2 virtual metres; wall trigger radius = 0.2 virtual metres), via a button press, which resulted in them turning green.

After all tokens had been collected, and all sensors activated, the participant completed a judgement of relative direction (JRD) criterion task, which was used to assess their knowledge of the VE’s layout (Figure 1C). On each trial of the JRD task, the participant was presented with a static picture of a global landmark (1s), which they were required to imagine facing. After a brief pause (0.5s), a picture of a different global landmark was shown and the participant was required to indicate the direction of the second landmark relative to the first. Specifically, if the participant thought that, when facing the first landmark, the second landmark was located to the participant’s left then they pressed the thumb button on the mouse (i.e., the left-most button); if the second landmark was located behind them, they pressed the left mouse button (i.e., the middle of the three response buttons), and if it was located to the right, they pressed the right mouse button (i.e., the right button). Performance was assessed via the number of correct responses, with the participant proceeding to the fMRI scanner task only if they answered all 12 JRD questions correctly; an incorrect response resulted in the participant returning to the VE to repeat the exploration phase.

### fMRI task

In the fMRI task, the participant viewed passive movement in the VE and was required to indicate the global landmark located in the direction of a cue object positioned either to the left or right of their path. Each trial comprised passive movement along a predefined path in which the participant could see 1) one global landmark towards which they were moving, 2) one boundary, and 3) the cue object (Figure 2A). After the movement ended (two seconds), the screen faded to black for four seconds before the start of the decision phase (two seconds), which comprised a forced-choice response. Here, the participant had to indicate which of the three global landmarks (i.e., the remaining landmarks not seen during the passive movement on the trial) was located in the direction of the cue object. For example, if the participant viewed a path heading towards the mountain, and the cue object was positioned on the right-hand side of the path, the participant was required to identify the global landmark located to the right of the mountain. In this case, the correct response would be the clock tower. In the forced-choice decision, the three global landmarks were presented on screen in a row, with the position of the landmarks randomly assigned either to the left, middle, or right position of the screen on each trial; randomising the screen position-landmark associations was important to ensure that they did not confound any subsequent decoding analyses (Supplementary Figure 1). The participant had to select, via a right-hand MR-compatible button box, which of the landmarks they thought was located in the direction of the cue object using either a thumb, index, or middle finger response, corresponding to the landmark image’s position on the screen.

**Figure 2.**
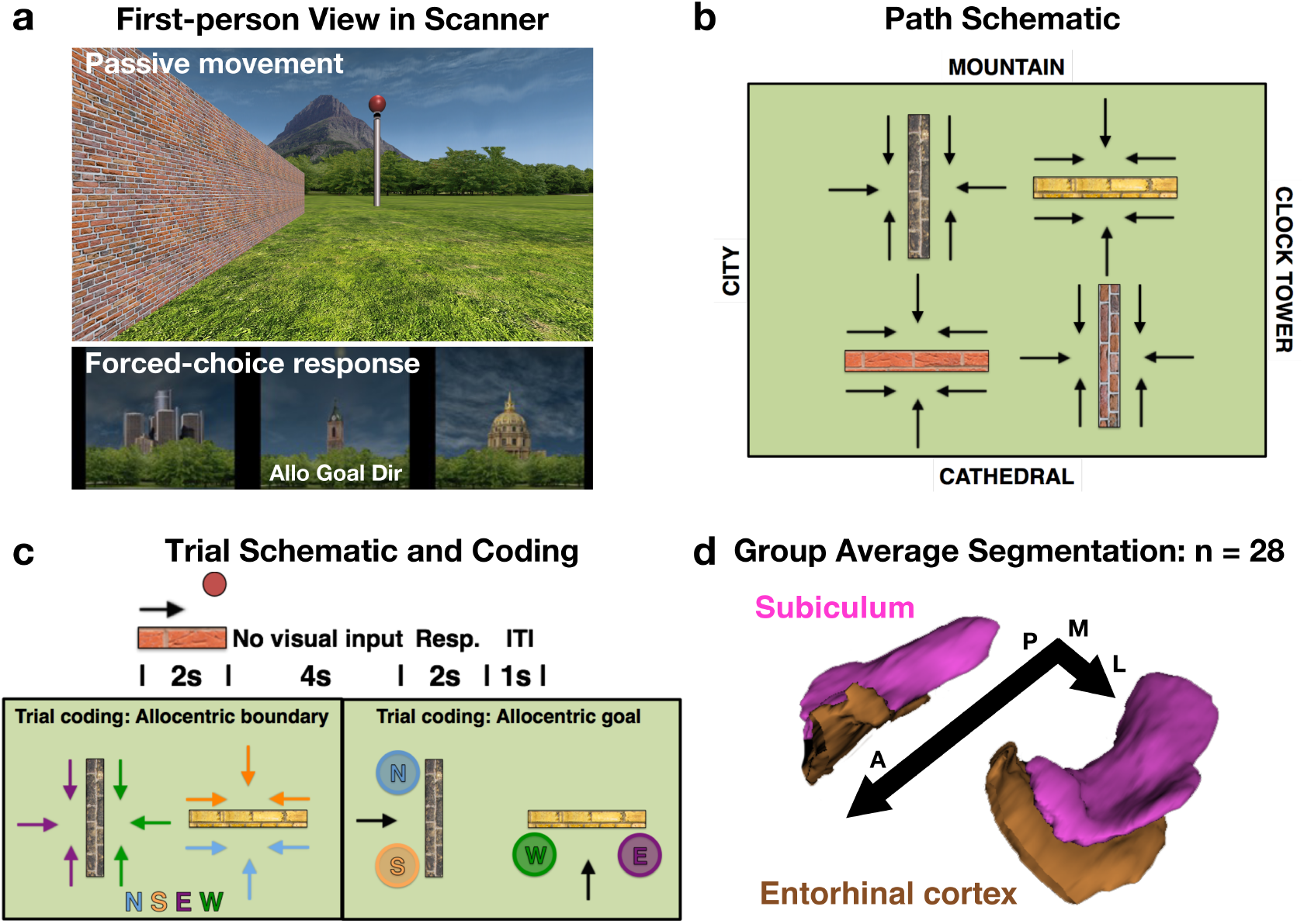
fMRI scanner task, trial coding, and example group ROIs. (a) On each scan trial, the participant viewed the global landmark towards which they were travelling, a boundary, and a cue object (pole with ball). The participant then completed a forced-choice decision in which they were presented with the three other global landmarks (i.e., those not seen during the passive movement) and were required to select the landmark located in the direction of the goal object (Correct answer here indicated by ‘Allo Goal Dir’). (b) Schematic of the 24 passive paths used in the scan task. Each path was repeated four times resulting in 96 trials per run. (c) Each trial comprised two seconds of passive movement, four seconds in which the visual input was removed (i.e., the portion of the trial used for the fMRI analysis), and two seconds to make the cue object direction decision. Trials were then coded according to either the allocentric boundary direction or the allocentric goal direction. (d) All analyses were carried out in the participant’s native EPI space using manually segmented masks of the entorhinal cortex and subiculum; the group averaged ROIs are presented here for display purposes only.

By using predefined paths in the VE we were able to control the position of the boundary and goal object relative to the participant. There were three paths per side of the boundary (24 in total), and these paths resulted in the boundary being located either to the left, right, or straight in front of the participant (see Figure 2B). Each path was repeated four times per run (96 trials per run; three runs in total) and the cue object’s position changed over trials so that its position was balanced across the left and right side of the path (i.e., for each path repeated four times per run, the cue object was located twice to the right, and twice to the left of the path). Trials could then be binned to examine different questions regarding allocentric boundary or allocentric goal direction coding (Figure 2C). Importantly, these different spatial properties were balanced across the different conditions, meaning that the comparisons were orthogonal. For example, trials used to examine allocentric boundary to the North would comprise views of two different boundaries (with their distinct textures), views of different global landmarks, the egocentric location of the boundary location to the participant’s left, right, and front, allocentric goal locations pointing to all four global landmarks, as well as an equal number trials in which the goal object was located egocentrically to the left or right of the participant. Each trial lasted 8s with a mean 1s inter-trial interval, and each of the three runs lasted 14.8 minutes.

### fMRI data acquisition

Imaging data were acquired using a 3T SIEMENS (Erlangen, Germany) Magnetom Prisma scanner, with a 64-channel phased array head coil. Scans comprised a whole-head, three-dimensional structural T1-weighted anatomical image with 1 mm isotropic resolution (TR/TE/Inversion time = 2500/2.82/1100 ms; flip-angle = 7 degrees; FOV = 256 × 256 mm^2^; 192 slices; GRAPPA acceleration factor 2); a high resolution moderately T2-weighted structural image comprising the hippocampus and EC acquired perpendicular to the long axis of the hippocampus using a turbo-spin-echo sequence (in-plane resolution = 0.4 × 0.4 mm^2^, slice-thickness = 1.5 mm; TR/TE = 4540ms/44ms; FOV = 224 × 224 mm^2^; 32 slices); gradient echo field maps (in-plane resolution = 1.6 × 1.6 mm^2^; slice-thickness = 2 mm; TR/TE1/TE2 = 720/4.92/7.38 ms; flip-angle = 60 degrees; FOV = 220 × 220 mm^2^; 72 slices), and three runs (445 volumes each) of T2*-weighted functional images acquired with a partial-volume echo-planar imaging sequence, aligned with the long axis of the hippocampus (in-plane resolution = 1.5 × 1.5 mm^2^, slice-thickness = 1.5 mm + 10% gap; TR/TE = 2000/30 ms; flip angle = 90 degrees; FOV = 192 × 192 mm^2^; 26 slices; GRAPPA acceleration factor 2).

### fMRI data preprocessing

A custom preprocessing pipeline was created using Nipype^27^, in which we combined packages from SPM12^28^, FSL5^29^, and Advanced Normalisation Tools^30^ (ANTS 2.1). The pipeline comprised realignment of the EPI data to the first volume of the series (SPM), intensity-normalization (FSL), and high-pass filtering with a cut-off of 128s (FSL). Structural T1 images were bias-corrected (SPM) and segmented (SPM), with the resulting grey matter, white matter and CSF tissue probability maps combined to create a mask for brain extraction.

Using FSL’s epireg, EPI data were coregistered to the structural T1, whilst also applying field map correction to the functional images using field maps acquired during scan sessions. High-resolution T2 images were coregistered to the T1 using ANTS. Manually segmented hippocampal T2 ROIs were coregistered to the EPI data by concatenating the T2-to-T1, and EPI-to-T1-inverse matrices using ANTS (Supplementary Figure 2). The EPI-to-T1-inverse matrix was used to move the T1 brain mask (comprising grey matter, white matter and CSF) into EPI space. The EPI data were then multiplied by this brain mask to remove all non-brain tissue.

### Medial temporal lobe masks

Bilateral hippocampi and parahippocampal cortices were segmented manually on the individual subjects’ T2 images using ‘ITK-SNAP’^31^, following an established protocol^32^ (Figure 2D). Given the differences in connectivity along the anterior and posterior portions of the EC and subiculum, we split each of the individual participant’s ROIs in the middle of the long axis, separately for each hemisphere. Due to movement artifacts in the T2 images, it was not possible to segment the hippocampi of three male participants. Consequently, all results reported below reflect the data from 28 participants (13 females).

### Data analysis

Prior to decoding analysis, movement parameters obtained from the realignment of the functional images were regressed out of the data. Here, we included 24 regressors in the model, reflecting the realignment parameters, their derivatives, their squares, and their square derivatives.

Each of the 96 trials per run was modelled separately in the analysis. To reduce the possible influence of visual information in our decoding analysis, we analysed the portion of data corresponding to the period of the trial after the passive movement ended during which there was no visual input (i.e., a black screen) and was therefore matched across different allocentric boundary/goal directions. To account for the lag of the haemodynamic response function, we analysed the volumes occurring 8 seconds (i.e., four volumes) after the onset of this period of the trial, and averaged the data over the next consecutive three volumes^33^. To enhance the signal corresponding to the allocentric condition of interest whilst maintaining the voxel space, we created an average over the three runs by first ordering the trials in each run according to the condition to-be-decoded (allocentric boundary or allocentric goal). The rationale here was to strengthen the condition of interest, whilst weakening any signal associated with other conditions (e.g., head direction). This trial-averaging resulted in 96 samples per participant, balanced equally across North, South, East and West directions for allocentric boundary and allocentric goal conditions.

A support vector classifier with L2 regularization as implemented in Scikit-learn^34^ was used for the decoding of different allocentric directions. The regularization strength was determined by adjusting the C hyperparameter. In order to follow decoding best practices^35^, we used nested cross-validation to estimate the best C hyperparameter and obtained a cross-validated estimate of the classifier accuracy with three outer folds using 20% of the data as test set in each fold. The best hyperparameter was chosen within the inner nested cross-validated fold, using a grid search with possible values in the range of 1 to 10^3^ in steps of power of 10. All decoding analyses were conducted in the participants’ native EPI space.

### Statistical tests

All behavioural data (mean accuracy and reaction time data from the learning phase JRD and fMRI task) were submitted to repeated-measures ANOVAs calculated using SPSS (IBM Corp. Version 21.0). Mauchly’s test of sphericity was used to assess homogeneity of variance for the ANOVAs, and Greenhouse-Geisser estimates of sphericity used to correct degrees of freedom when this assumption was violated. Given that we had no apriori predictions as to differences in performance across the different conditions of the behavioural tasks, follow-up paired sample T-tests interrogating significant main effects and/or interactions were Bonferroni-corrected for multiple comparisons. Effect sizes were calculated using online tools^36^, and all plots were created using a combination of Matplotlib^37^ and Seaborn^38^.

For the decoding analyses in the separate ROIs, we obtained the mean decoding score per participant over the three-folds of the cross-validation. We then used the bias-corrected and accelerated boot-strap^39^ (BCa) to sample from these values 10,000 times to obtain the distribution of our group-level decoding accuracy^40^. Non-parametric methods were used to generate a p-value based on the distribution of our data, where we first subtracted the group mean decoding accuracy from each participant’s decoding score, before adding chance performance (i.e., 25%). This had the effect of shifting the distribution of our group’s decoding scores to around chance performance, and we then again used the BCa (with 10,000 samples) with these values to generate our null distribution. The one-tailed p-value was calculated by counting the number of times the boot-strap null mean exceeded our observed group-level decoding score and dividing this value by the number of samples (i.e., 10,000). Outside of our key ROIs (EC and subiculum) we tested also whether we could decode allocentric boundary and goal direction in manually segmented masks of the CA1, CA23DG, PHC. Given that we did not have overt predictions as to the expected pattern of results in these ROIs, we used a Bonferroni-adjusted alpha level to test for significant effects (three ROIs × two conditions = 0.05 / 6 = 0.008 adjusted alpha).

To test directly whether there was evidence of an anterior-posterior difference in decoding accuracy according to task in the EC and subiculum, we submitted participants’ mean decoding scores to a repeated-measures ANOVA comprising the factors i) ROI (EC, subiculum) ii) Anterior/posterior section, and iii) Condition (allocentric boundary direction, allocentric goal direction). Given our apriori predictions as to anterior versus posterior differences in decoding accuracy according to allocentric condition, follow-up two-tailed t-tests were not corrected for multiple comparisons.

## Results

### Behavioural

Before proceeding to the scanned fMRI task, participants were required to learn the layout of the VE as assessed by a JRD task. Participants needed on average 3.39 (SD = 1.01) rounds of exploration to learn the relative directions of the global landmarks. To test whether performance on the JRD changed as a function of the initial landmark facing direction, or the angular disparity of the second landmark relative to the first, accuracy and reaction time (RT) data were submitted to separate repeated-measures ANOVAs comprising the factors Landmark (Mountain, Cathedral, Clock tower, City) and Angular Disparity (90, 180, 270 degrees). Accuracy was modulated by the angular disparity of the second landmark [F (2, 54) = 9.49, p = 0.0003, ω_*p*_^2^ = 0.230], but not the initial landmark facing direction [F (3, 81) = 0.41, p = 0.74, ω_*p*_^2^ = −0.021], and these two factors did not interact [F (6, 162) = 0.34, p = 0.92, ω_*p*_^2^ = −0.023] (Figure 3A). Performance was significantly better for landmarks located at 180, or 90 degrees disparity versus those located at 270 degrees disparity ([t (27) = 3.45, p = 0.001, 95% CI [0.05, 0.18], Hedges’s *g_av_* = 0.84; adjusted alpha = 0.05 / 3 comparisons = 0.017], and [t (27) = 3.66, p = 0.001, 95% CI [0.04, 0.14], Hedges’s *g_av_* = 0.73], respectively); performance did not differ between 180 and 90 degree disparities [t (27) = 0.93, p = 0.36 95% CI [-0.07, 0.03], Hedges’s *g_av_* = 0.18]. The same analysis for RT data revealed a similar pattern of results, with a main effect of angular disparity [F (1.16, 31.36) = 9.37, p = 0.003, ω_*p*_^2^ = 0.224], but no effect of initial landmark facing direction [F (1.44, 38.98) = 0.73, p = 0.54, ω_*p*_^2^ = −0.009], and no interaction between these factors [F (6, 162) = 0.61, p = 0.72, ω_*p*_^2^ = −0.013]. Responses were fastest for landmarks located at 180 degrees disparity versus those located at 270 [t (27) = 3.27, p = 0.003, 95% CI [−4.29, −0.98], Hedges’s *g_av_* = 0.70; adjusted alpha = 0.05 / 3 comparisons = 0.017], and a trend towards being faster than those located at 90 degrees [t (27) = 2.4, p = 0.02, 95% CI [−2.7, −0.21], Hedges’s *g_av_* = 0.48]; responses for landmarks located at 90 degree disparity were faster than those located at 270 degrees [t (27) = 3.71, p = 0.0009, 95% CI [−1.83, −0.53], Hedges’s *g_av_* = 0.26].

**Figure 3.**
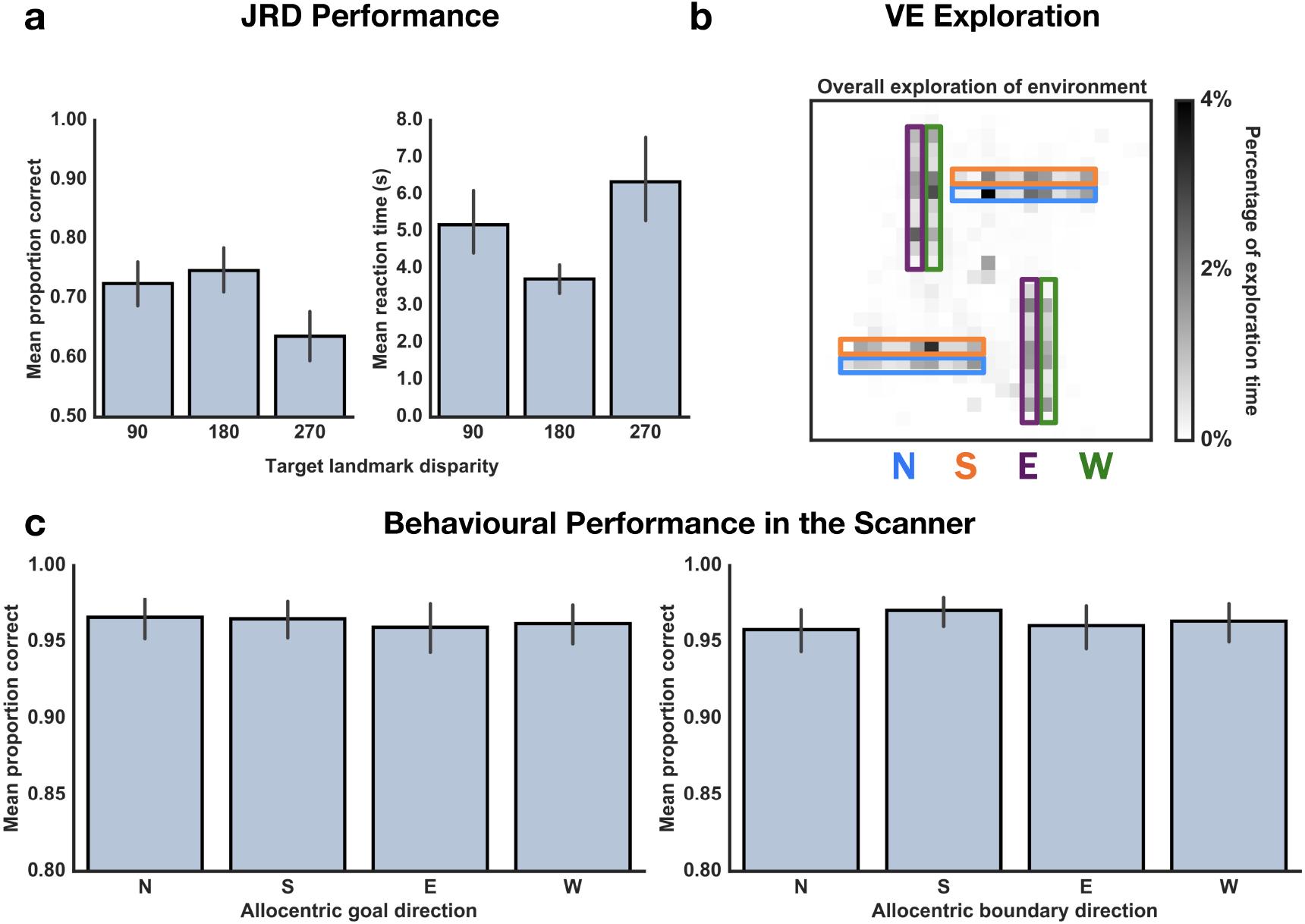
Mean performance during the learning phase and fMRI scan task. (a) On the JRD task, participants made more accurate judgements regarding landmarks located either at 180, or 90 degrees angular disparity, and made the fastest responses for landmarks located at 180 degrees disparity. (b) Exploration time in the environment was matched across the four different allocentric boundary directions. (c) In the fMRI scan task, accuracy was matched for allocentric goal direction, and the same was true when grouping the trials according to allocentric boundary direction. Error bars represent the 95% CI.

Participants were required to understand that the VE boundaries were impassable. It was crucial, therefore, that they spent a large proportion of their exploration time in close proximity to the boundaries. To assess time spent near to the boundaries, for each participant we divided the total explorable space of the VE into 10,000 bins (each bin = 5 × 5 virtual metres^2^ of the VE), and normalised the total time across all rounds of exploration so that it summed to 1. We then created masks of the areas next to the boundaries in the VE, comprising the four different allocentric boundary directions (each mask per boundary side was 40 × 5 virtual metres). On average, participants spent 76.16% (SD = 10.45%) near the environment boundaries (Figure 3B). A one-way repeated-measures ANOVA revealed that exploration time did not differ as a function of allocentric boundary direction [F (2.27, 61.24) = 1.16, p = 0.33, ω_*p*_^2^ = 0.006].

Given the extensive training required prior to fMRI scanning, performance on the scanner task was very high (Figure 3C). Accuracy and RT data were submitted to separate one-way repeated-measures ANOVAs comprising Allocentric Goal Direction (North, South, East, West). Accuracy was matched across all four allocentric goal directions [F (2.18, 58.89) = 0.35, p = 0.73, ω_*p*_^2^ = −0.023], whereas RT differed [F (3, 81) = 29.2, p = 0.00001, ω_*p*_^2^ = 0.499]. Follow-up comparisons revealed that responses to allocentric goals located to the North were significantly quicker than those goals located to the South [t (27) = 3.15, p = 0.004, 95% CI [−0.06, −0.01], Hedges’s *g_av_* = 0.36; adjusted alpha = 0.05 / 6 comparisons = 0.008], East [t (27) = 6.72, p < 0.001, 95% CI [−0.09, −0.05], Hedges’s *g_av_* = 0.65], and West [t (27) = 10.20, p < 0.001, 95% CI [−0.12, −0.08], Hedges’s *g_av_* = 0.85]. Similarly, responses to goals located to the South were faster than those located to the East [t (27) = 3.12, p = 0.004, 95% CI [−0.05, −0.01], Hedges’s *gav* = 0.3] and West [t (27) = 5.06, p = 0.00003, 95% CI [−0.09, −0.04], Hedges’s *g_av_* = 0.54]; differences in reaction times for responses to goals located to the East and West did not survive Bonferroni-correction [t (27) = 2.45, p = 0.021, 95% CI [−0.06, −0.01], Hedges’s *g_av_* = 0.26]. Although the participants performed a task regarding the allocentric goal direction, trials could be coded also according to the allocentric boundary direction. The same repeated-measures ANOVAs were conducted with this coding, and again revealed that accuracy was matched across allocentric boundary direction [F (3, 81) = 1.87, p = 0.14, ω_*p*_^2^ = 0.03], but that reaction time differed [F (3, 81) = 8.24, p = 0.00007, ω_*p*_^2^ = 0.204]. The differences in reaction time stemmed from faster responses to trials in which the allocentric boundary was located to the North relative to East [t (27) = 3.34, p = 0.002, 95% CI [−0.04, −0.01], Hedges’s *g_av_* = 0.23; adjusted alpha = 0.05 / 6 comparisons = 0.008], and West [t (27) = 2.91, p = 0.007, 95% CI [−0.04, −0.01], Hedges’s *g_av_* = 0.19]. The same pattern of data was evident for allocentric boundaries located to the South, with faster responses relative to East [t (27) = 4.08, p = 0.0003, 95% CI [−0.05, −0.02], Hedges’s *g_av_* = 0.28] and West [t (27) = 3.45, p = 0.002, 95% CI [−0.04, −0.01], Hedges’s *g_av_* = 0.24]; reaction times did not differ for allocentric boundaries located to the North and South [t (27) = 0.75, p = 0.46, 95% CI [−0.01, 0.02], Hedges’s *g_av_* = 0.06], nor East and West [t (27) = 0.69, p = 0.49, 95% CI [-0.01, 0.01], Hedges’s *g_av_* = 0.04] (Supplementary Figure 4).

### Multivariate decoding

Mean group-level decoding accuracy of allocentric boundary direction in the posterior EC was significantly above chance (Figure 4A), as determined by the BCa null distribution analysis, where none of the bootstrap null group mean decoding accuracies exceeded the observed group decoding accuracy [p = 0.0]. The same analysis for allocentric goal direction, however, did not exceed chance performance [p = 0.515]. In the anterior EC (Figure 4B), it was possible to decode also allocentric boundary direction [p = 0.024], however in contrast to the posterior EC, the anterior EC contained information also regarding the allocentric goal direction [p = 0.002].

**Figure 4.**
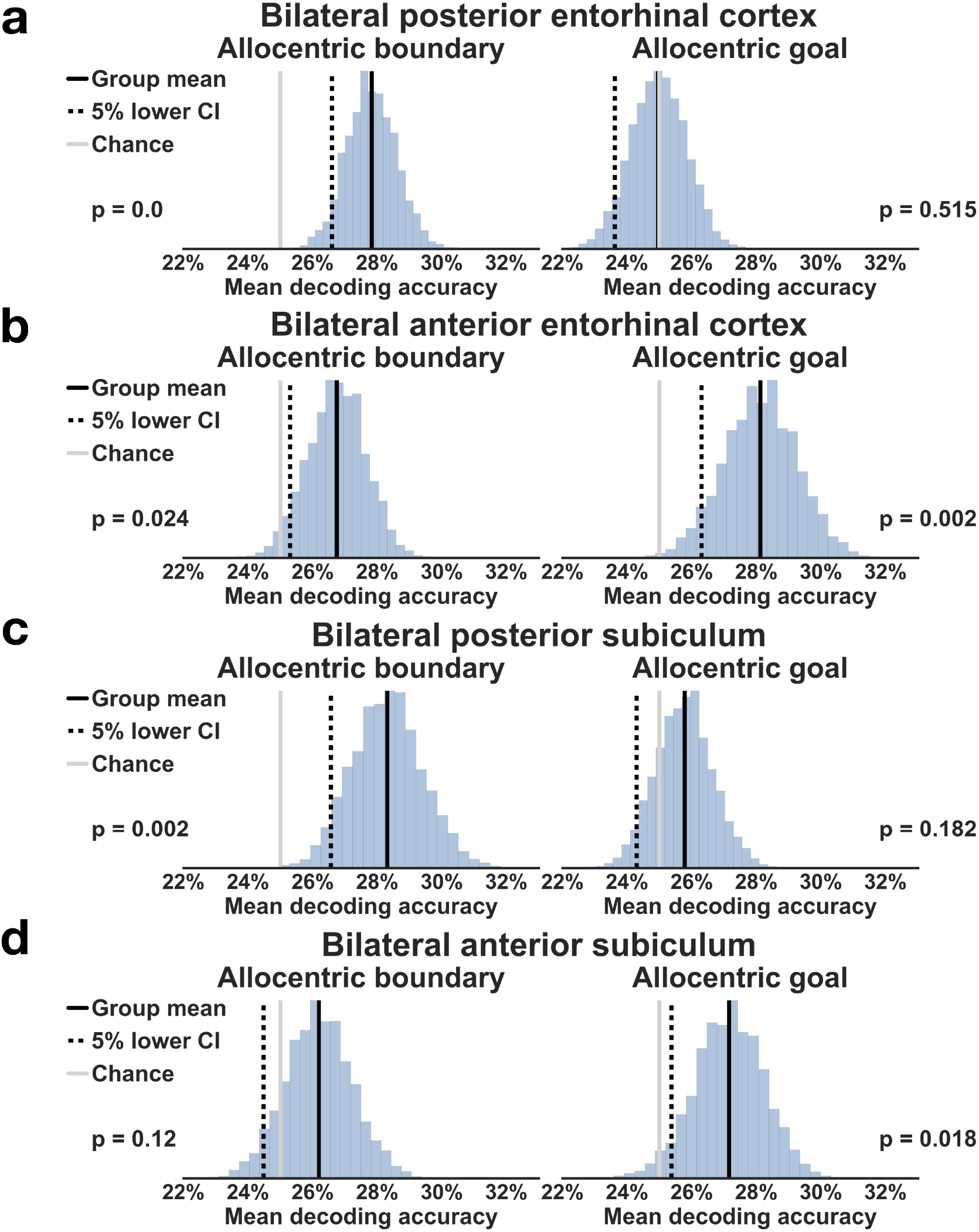
fMRI decoding results in bilateral entorhinal cortex and subiculum. (a) Using a linear SVC with L2 regularisation, in bilateral posterior EC group decoding accuracy for allocentric boundary direction was significantly above chance, as determined by a boot-strap permutation test (10,000 samples). It was not possible, however, to decode allocentric goal direction. (b) In anterior EC, we were able to decode both allocentric boundary and goal direction. (c) The posterior subiculum contained similar information to the posterior EC, in that we could decode allocentric boundary, but not goal, direction. (d) Finally, in anterior subiculum it was possible to decode allocentric goal, but not boundary, direction.

The pattern of data in the posterior subiculum mirrored that of the posterior EC, with significantly above chance decoding of allocentric boundary direction [p = 0.002], but not allocentric goal direction [p = 0.182] (Figure 4C). In line with anterior EC, the anterior subiculum contained information regarding the allocentric goal direction [p = 0.018] (Figure 4D). It was not possible in this region, however, to decode allocentric boundary direction [p = 0.12].

To test directly for an anterior-posterior difference in decoding accuracy according to task, we entered participants’ decoding scores into a repeated-measures ANOVA, comprising the factors ROI (EC, subiculum) × Section (Anterior, posterior) × Condition (Allocentric boundary, allocentric goal). The ANOVA confirmed that decoding accuracy was modulated by the anterior versus posterior portion of the ROI, for different tasks [F (1, 27) = 10.85, p = 0.003, ω_*p*_^2^ = 0.253] (Figure 5), but this effect was not modulated by ROI as evidenced by the absence of a significant three-way interaction [F (1, 27) = 0.06, p = 0.81, ω_*p*_^2^ = −0.034]. Follow-up t-tests interrogating the Section × Condition interaction revealed a double-dissociation such that decoding accuracy was higher for allocentric boundary in posterior versus anterior regions [t (27) = 2.58, p = 0.015, 95% CI [0.33, 2.87], Hedges’s *g_av_* = 0.56], whereas the reverse pattern of data was true for allocentric goal decoding with greater accuracy for anterior versus posterior regions [t (27) = 2.39, p = 0.024, 95% CI [0.32, 4.22], Hedges’s *g_av_* = 0.61]. In posterior sections of the ROIs, decoding accuracy for allocentric boundary was also significantly greater than allocentric goal [t (27) = 3.02, p = 0.005, 95% CI [0.87, 4.53], Hedges’s *g_av_* = 0.89], but there was no difference in decoding accuracy between conditions in anterior sections of the EC and subiculum [t (27) = 1.35, p = 0.19, 95% CI [-0.60, 2.95], Hedges’s *g_av_* = 0.33].

**Figure 5.**
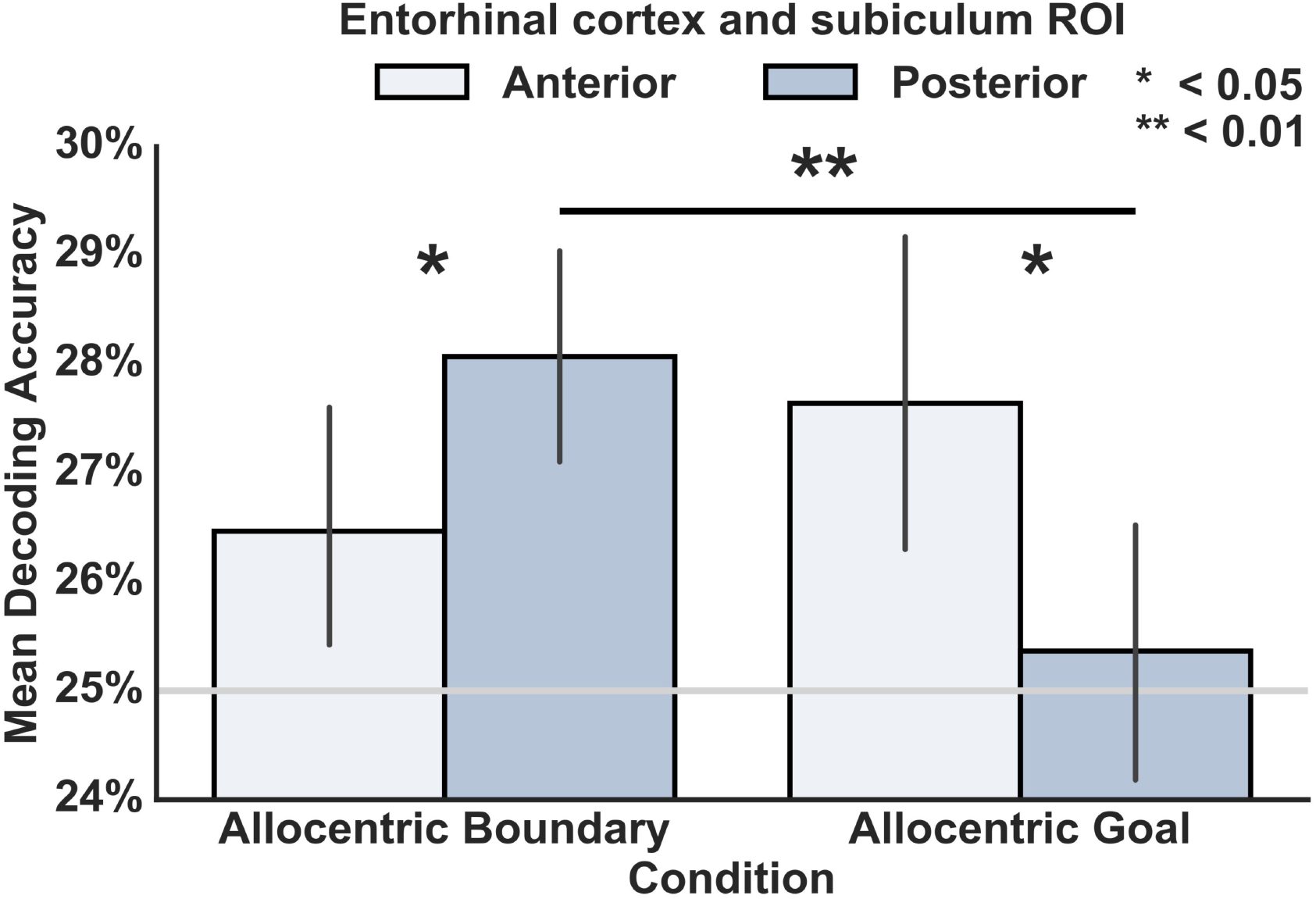
Testing for an anterior-posterior difference in decoding accuracy in the EC and subiculum according to task. The group’s decoding accuracies for allocentric boundary direction and allocentric goal direction were submitted to a ROI × Section × Condition ANOVA, which revealed a significant Section × Condition interaction. The interaction was driven by a double-dissociation in which posterior portions of the ROIs had greater mean decoding accuracies for boundary versus goal direction, whereas the reverse pattern of data was true in anterior regions. Error bars represent the 95% CI; the grey horizontal line represents chance decoding performance.

Outside of our EC and subiculum ROIs, we were able to decode allocentric goal direction in the PHC [p = 0.006]; decoding of allocentric boundary and goal direction did not survive Bonferroni-correction in either the CA1, or CA23DG (Supplementary Figure 5).

## Discussion

In the current study, we provide the first evidence that brain regions, analogous anatomically to those in the rodent brain, the posterior EC and subiculum, code for the allocentric direction to environment boundaries. Moreover, we found that anterior sections of these structures code for the allocentric goal direction. These data support the notion of a division of labour in the EC, in which different regions support processes involved in spatial navigation, and the coding of external sensory information^8^. Our findings are broadly consistent also with previous research in humans that has shown functional differences in the EC according to stimulus type (i.e., scenes versus objects)^6,7^.

Environment boundaries support successful navigation by providing an error correction signal when navigation is based upon path integration^12^, and static positional information during landmark navigation^22^. Consistent with the rodent literature, we were able to decode allocentric boundary direction in posterior EC and subiculum. Although previous studies have shown univariate responses associated with boundaries^23,24^, by using multivariate analysis methods, we have provided the first evidence that this medial temporal lobe boundary signal contains also the allocentric information crucial for both grid and place cell function. Moreover, in contrast to previous research, we were able to separate the contribution of allocentric boundary and allocentric goal coding^25^. We observed above chance decoding of allocentric boundary direction also in anterior EC, albeit with significantly lower accuracy than that observed in posterior EC^41,42^. Although border cells are more prominent in the rodent MEC versus LEC, our data is in line with a weak boundary signal that has been reported in rodent LEC^42^, and may be explained by the transmission of information between the two regions, due to their high levels of connectivity. Given that we see the same pattern of data in posterior EC and subiculum, it might lead to questions as to whether this represents a redundancy of function, with the same information represented in both regions. One possible difference between the posterior EC and subiculum might be information regarding the distance to the environment boundary. Although we manipulated only allocentric boundary direction in the current study, a key component for boundary vector cells is that they also code for distance to a boundary^18^, with evidence of this coding coming from recordings from the subiculum of intracranial implant patients^24^. Examining the sensitivity of different brain regions to boundary proximity remains an important future question for boundary coding research, and the subiculum would be a likely candidate to contain this information.

In contrast to MEC, the rodent LEC receives direct projections from the object-sensitive perirhinal cortex, and it has been hypothesised that it codes for external sensory information, such as prominent objects in specific locations that may constitute landmarks^8^. Consistent with this interpretation, we observed above chance decoding of allocentric goal direction in anterior EC and subiculum. These data support previous studies in which increased activity in anterior hippocampus is associated with successfully navigating to a goal location^43^. Evidence from rodent studies suggests that not only does the rodent LEC code for objects in specific places, but that it is more responsive to local cues rather than distal landmarks^44^. Specifically, when two sets of cues (local versus distal) were placed in opposition, the population response of LEC neurons tracked changes to the local cues. In the current study we did not manipulate global versus local features, but it is conceivable that the “global versus local’ division of labour in EC emerges when there are multiple reference frames that need to be coordinated. Future studies in humans will be required to test whether the EC differentially codes for these different spatial cues.

Previous studies of EC function in humans have supported a division of labour according to object versus scene/spatial stimuli^6,7^. Why this distinction emerges according to stimulus type, however, remains unclear. One possibility is that, due to foveal vision, primates visually explore space, which in turn engages neural populations that support spatial navigation. For example, grid cell activity has been demonstrated during the passive viewing of scene stimuli^45^, with evidence also of neurons with a profile similar to that of border cells. Similar data have been reported recently in human fMRI whilst participants visually explored 2D scenes^46,47^. The scene versus object distinction, therefore, may reflect the engagement of these spatially-tuned neural populations during the viewing of scene stimuli. The current study helps elucidate further the exact mechanism underlying this stimulus-specific effect in the EC, and suggests that, in part, the processing of boundary information in scenes drives this scene-sensitive effect in posterior EC. In contrast, recent human studies demonstrate that objects are more likely to engage the anterior EC, which is closely connected with perirhinal cortex. The current data are not inconsistent with this notion, as the allocentric goal direction signal could be interpreted as reflecting the participant bringing to mind a specific landmark object. Understanding the precise role of the anterior EC, and what manipulations govern its involvement during spatial tasks, remains an important clinical objective, given, for example, that proteins such as tau aggregate in the transentorhinal area (comprising perirhinal cortex and anterior EC) early in Alzheimer’s disease^48^.

Neuronal populations originally thought to support only spatial navigation have been shown to be involved also in more abstract cognitive processes. For example, in rodents MEC grid cells map not only space, but also different sound frequencies^49^. Similarly, grid cell-like representations in humans, revealed via fMRI, have been found during spatial navigation^50^,imagined navigation^51^, and most recently in the organisation of conceptual knowledge^52^. In the current experiment, therefore, although we have used a very concrete example of a boundary coding (i.e., a physical boundary), the same posterior EC mechanism may support more abstract boundary-related processes, such as the segmentation of temporal information into event episodes, or the coding of visual boundaries^53^. There is evidence that boundaries are used to segment a continuous temporal stream into distinct episodic events. For example, increased forgetting of object pairs is observed when having to remember items between-rooms versus within the same room^54^. Furthermore, activity in the hippocampus has been shown to correlate with the salience of event boundaries during the viewing of films^55^. The ERC and subiculum, therefore, may also play a critical role in the formation of event episodes.

Outside of our key regions of interest, we observed significant decoding of allocentric goal direction in the PHC. The PHC has been shown to be exquisitely sensitive to scene stimuli, and in particular the structure of a scene^56,57^. Consistent with our findings, there is no evidence that the PHC codes for the allocentric direction to environment boundaries. There is data, however, to suggest that the PHC contains information regarding the direction to an imagined goal^58^. Specifically, when participants were required to remember the direction between two well-learned goal locations only the PHC was sensitive to similarities in imagined direction. These data are compatible with the findings reported here for a task in which participants were required to remember the allocentric direction to a goal landmark given the current heading direction. Alternatively, this above chance decoding could reflect the participants bringing to mind the specific landmark, with landmarks also known to engage the PHC^56^.

Taken together, our study provides the first evidence of allocentric boundary coding in humans, and suggests that, consistent with models of anatomical connectivity, posterior EC and subiculum provide support for positional coding, whereas the anterior EC and subiculum code for external sensory information such as landmarks. These findings advance our understanding of EC function, and provide further mechanistic explanation underlying the division of labour in this region.

## Supplementary Information

During the fMRI scanner task, the on-screen positions of the landmarks in the forced-choice response were randomly assigned (either left, middle, or right). This was important for the decoding analysis, so that no other information confounded the spatial property that we were trying to decode. To ensure that there was no systematic bias in the position of these items, for each landmark we calculated the proportion of times it was located in the left, middle, or right position on the screen (Supplementary Figure 1). These values were submitted to a repeated-measures ANOVA, comprising the factors Landmark (Mountain, Cathedral, Clock tower, City) × fMRI run (1, 2, 3) × Position (left, middle, right) and revealed that there were no significant main effects or interactions (all Fs < 1.44, ps > 0.20) relating to the spatial position of the items on screen.

For the decoding analyses, we used anatomical ROIs of the hippocampal subregions, comprising EC, subiculum, CA1, CA23DG, and PHC traced manually on individual participant’s T2-weighted images. Advanced Normalisation Tools was then used to move these anatomical ROIs to EPI space, using a composition of the EPI-to-T1 inverse, and T2-to-T1, matrices (Supplementary Figure 2). We then divided the EC and subiculum ROIs into anterior and posterior portions by dividing them at the midpoint along their longitudinal axis. In total we had seven bilateral ROIs, in which we assessed also the mean temporal signal-to-noise ratio (calculated by dividing each voxel’s mean intensity per run by its standard deviation over time, and averaging the resulting values across each ROI) and volume (Supplementary Figure 3).

Outside of our key ROIs (the EC and subiculum), it was possible to decode allocentric goal and allocentric boundary direction in the PHC (Supplementary Figure 5); decoding accuracies in the CA1 and CA23DG did not survive Bonferroni correction (p = 0.006).

**Supplementary Figure 1.**
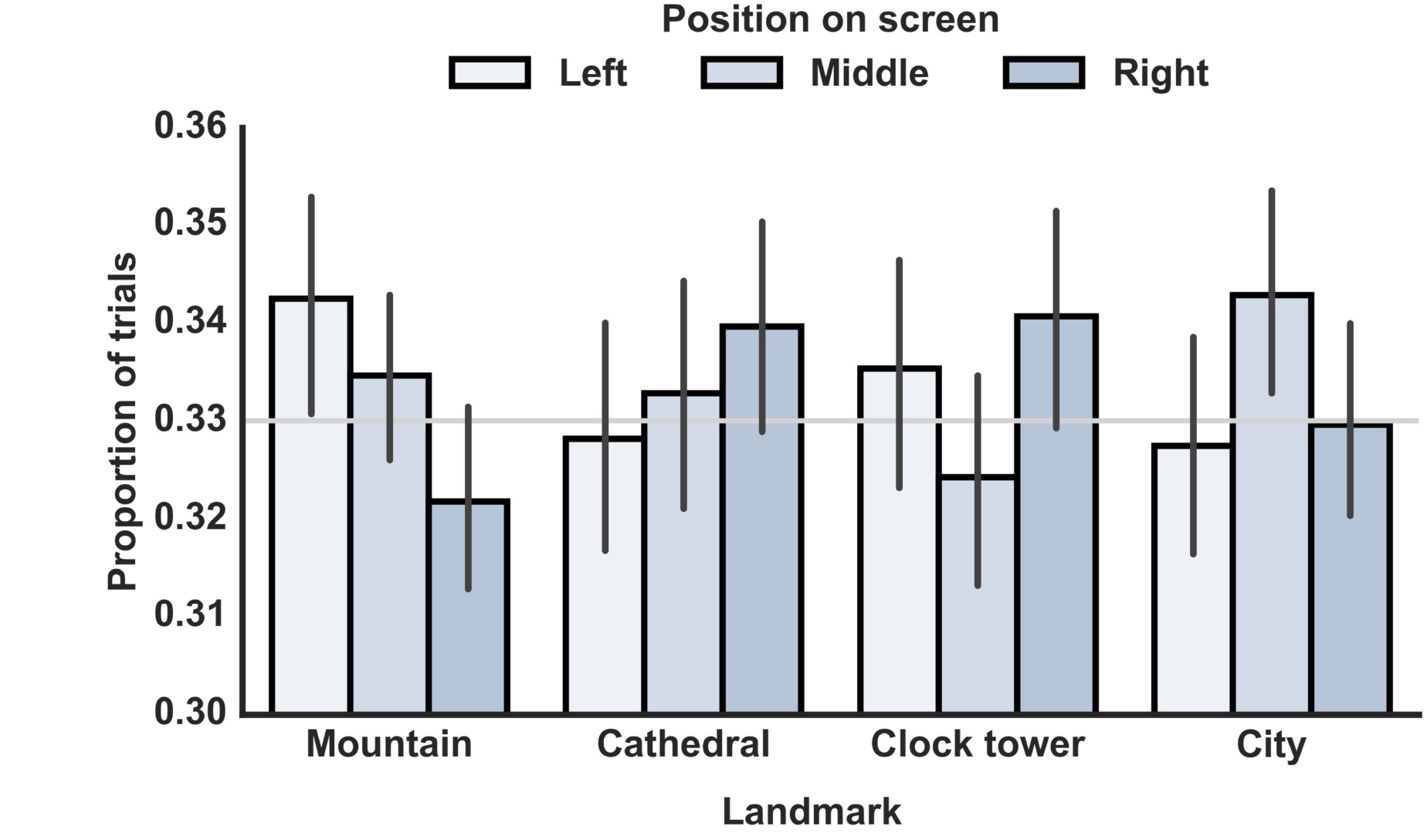
Proportion of trials in which each landmark was located either to the left, middle, or right of the screen during the forced-choice response in the fMRI task (see Figure 2a in the main text). To ensure that the position of the landmark on the screen (which corresponded also to the position on the button box) did not confound our decoding results, the position of the landmarks on screen was randomised, meaning that there was no systematic relationship between landmark and screen position. Error bars represent 95% CI; the grey horizontal line represents chance.

**Supplementary Figure 2.**
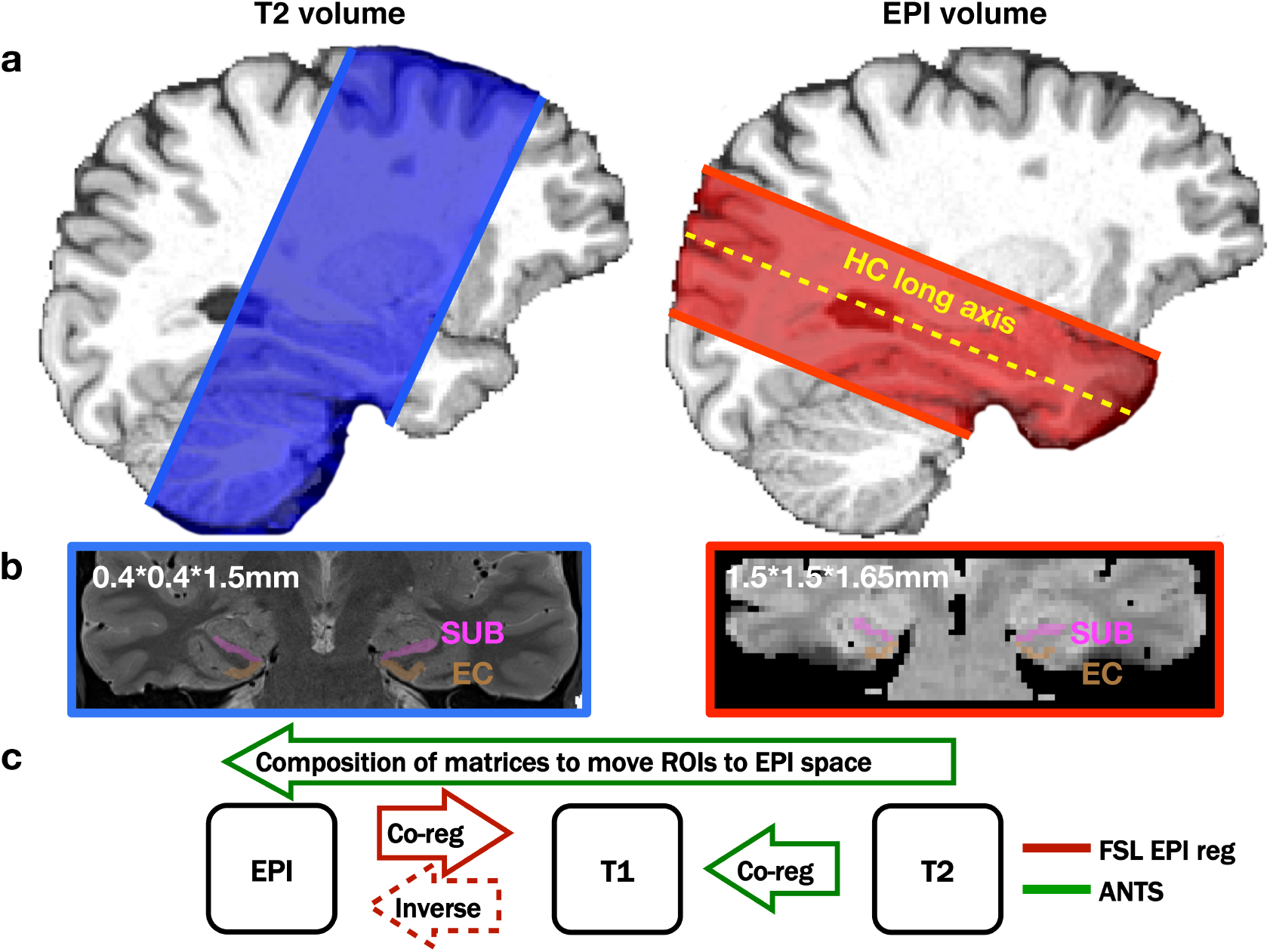
Co-registration of T2 and EPI volumes. (a) Bilateral hippocampi were segmented manually on individual subject’s T2-weighted images. (b) These masks were then co-registered with the high-resolution EPI slab aligned with the longitudinal axis of the hippocampus, using (c) a combination of FSL and ANTs.

**Supplementary Figure 3.**
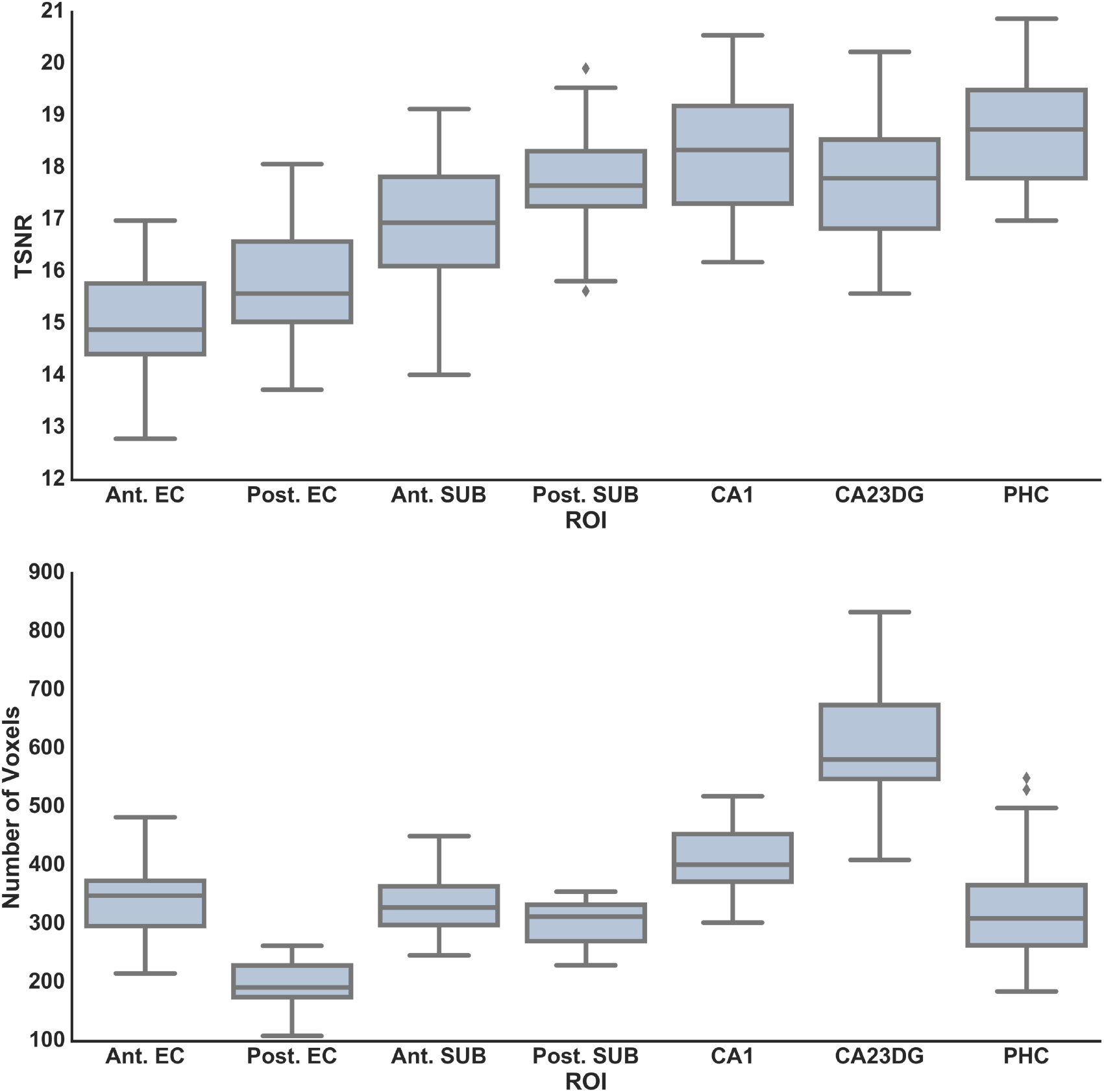
Temporal signal-to-noise ratio and number of voxels for each ROI, averaged over the group (n=28).

**Supplementary Figure 4.**
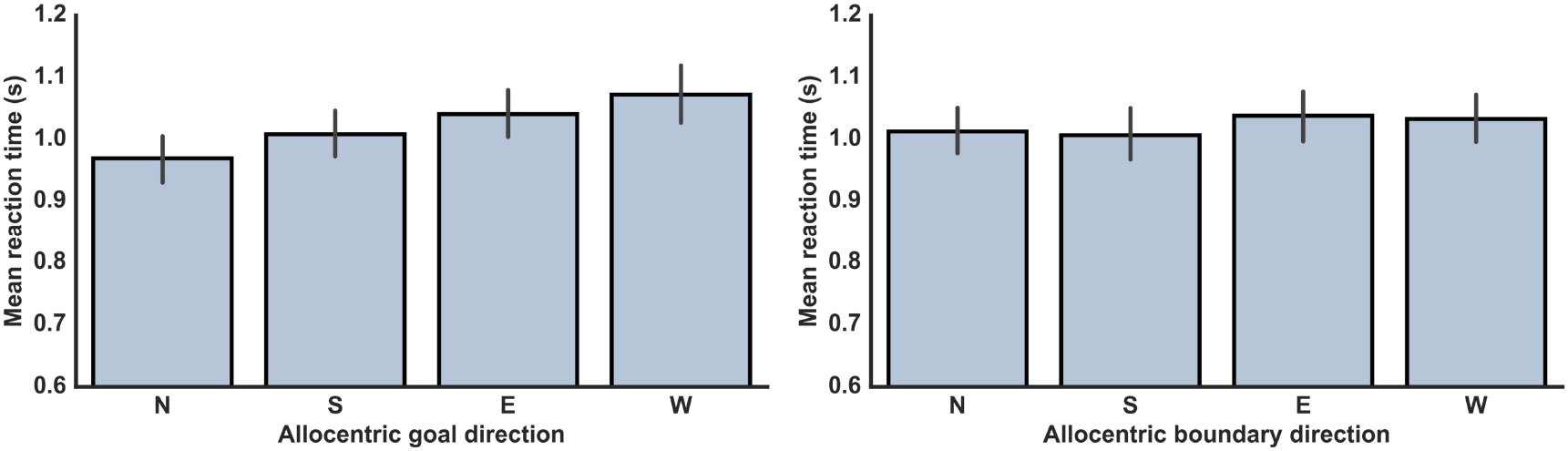
Reaction times on the fMRI scanner task with trials coded separately according to allocentric goal and allocentric boundary direction.

**Supplementary Figure 5.**
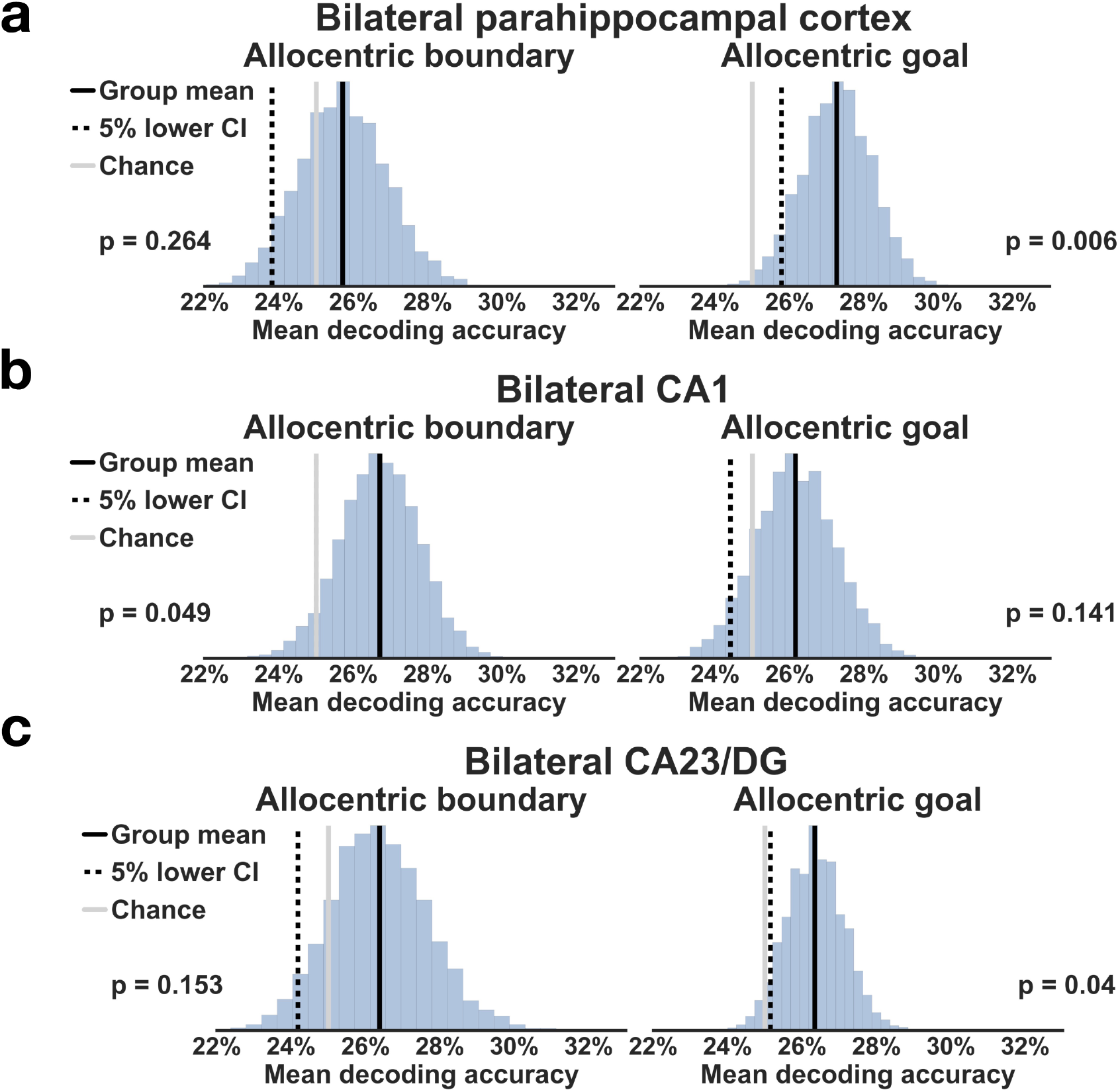
Decoding of allocentric boundary and goal direction outside the EC and subiculum. (a) In the PHC, it was possible to decode allocentric goal direction, (b) but group decoding accuracies for allocentric boundary and goal direction in bilateral CA1, and (c) CA23DG did not survive Bonferroni-correction (p = 0.008).

